# A spontaneous mutation in ADIPOR1 causes retinal degeneration in mice

**DOI:** 10.1101/2024.06.06.597783

**Authors:** Junzheng Yang, Natasha M. Buchanan, Erika Lima, Angela Banks, Valentin M. Sluch, Lin Fan, Barrett Leehy, Ivana Arellano, Yubin Qiu, Garrett Klokman, Shawn Hanks, Joanna Vrouvlianis, Vanessa Davis, Chung-Yeh Wu, Aaron Danilack, Dennis S. Rice

## Abstract

Adiponectin receptor 1 (ADIPOR1) is a transmembrane protein necessary for normal anatomy and physiology in the retina. In a recent study of complement factor H knockout mice (*Cfh*^−/−^), our lab discovered a flecked retina phenotype and retinal thinning by fundus imaging and optical coherence tomography (OCT), respectively. The phenotype was observed in a subset (50%) of *Cfh*^−/−^ mice. The thinning observed *in vivo* is due to an early degeneration of rod photoreceptors. This phenotype has not been reported in published studies of *Cfh*^−/−^ mice. AdipoR1 knockout mice (*AdipoR1*^−/−^) and mice deficient in Membrane Frizzled Related Protein (MFRP) exhibit this phenotype, suggesting an involvement in the emergence of the retinal degeneration observed in a subset of *Cfh*^−/−^ mice. *Cfh* and *AdipoR1* are located in close proximity on mouse Chromosome 1 (Chr1) and a complementation cross between *Cfh* and *AdipoR1* mice with retinal degeneration produced 100% progeny with retinal degeneration. Sequencing of the *Cfh*−/− mice revealed a c.841 C > T mutation in *AdipoR1*. Furthermore, one *Cfh* wildtype (of Cfh^+/+^) and 2 heterozygous (of *Cfh*^+/−^) mice exhibited retinal degeneration and were homozygous for the point mutation. The c.841 C > T mutation results in a proline to serine conversion at position 281 (P281S) in ADIPOR1. This residue is critical for ADIPOR1 open and closed conformations in the membrane. *In silico* modeling of candidate ADIPOR1 ligands, 11-cis-retinaldehyde and docosahexaenoic acid (DHA), that are deficient in AdipoR1^−/−^, suggests that ADIPOR1 is involved in trafficking retinoids and fatty acids and their combined deficiency in the ADIPOR1 mutant retinas might explain the retinal degeneration phenotype.

## Introduction

Age-related macular degeneration (AMD) is a medical condition that primarily impacts the macula, a crucial part of the retina in the eye. This condition is a leading cause of global vision loss, accounting for about 8.7% of all cases of blindness. AMD is common in developed countries and tends to primarily affect people over the age of 60. A mix of genetic and environmental factors influence the development and progression of AMD. Key processes in the body, such as the immune system’s response (complement system), fat metabolism (lipid pathways), blood vessel formation (angiogenic pathways), inflammation, and the health of the extracellular matrix, all play a role in the development of this disease [1].

Recent advancements in understanding age-related macular degeneration have been significantly driven by research into the role of the complement system [2]. Key among these advancements is the identification of a connection between AMD and a widespread single-nucleotide polymorphism (SNP) in the gene for complement factor H (CFH). Factor H is crucial in regulating the alternative pathway of the complement system, playing a pivotal role in safeguarding host cells and tissues against collateral damage during complement activation.

We acquired *Cfh*-deficient mice from Pickering and Botto [3] to further study the role of complement biology in the eye. The founder line displayed a normal fundus phenotype and marginal thinning of the outer nuclear layer in aged mice. Subsequent breeding through multiple generations produced a subset of *Cfh*^−/−^ mice that presented with early-onset retinal degeneration associated with a uniform, flecked retina fundus appearance. This phenotype consistently appeared in a subgroup of *Cfh*^−/−^ mice, suggesting that the Cfh deficiency itself does not cause the observed retinal lesions. Furthermore, the retinal phenotypes in the subset of *Cfh*^−/−^ mice phenocopies those observed in *Adipor1^−/−^* mice [4].

The importance of ADIPOR1 in retinal health is conserved across species, suggesting a critical biological function. Mutations in human ADIPOR1 are associated with retinitis pigmentosa, morpholino knockdown in Zebrafish AdipoR1 reduced the number of photoreceptors [5] and knockout mice have an early onset a progressive loss of photoreceptors. A SNP in human ADIPOR1 is associated with increased risk of age-related macular degeneration in a Finnish population. Here we describe a novel and important spontaneous mutation in murine AdipoR1 that will facilitate future studies on AdipoR1 function and informs a hypothesis for the role of AdipoR1 in the transport of polyunsaturated fatty acids and retinoids in the eye.

## MATERIALS AND METHODS

### Animals

The animal work in this paper adheres to the ARVO Statement for the Use of Animals in Ophthalmic and Vision Research and was approved by the NIBR IACUC Committee. Intercross breeding of *Cfh^+/−^* mice from the current line was used to generate study cohorts which ranged in age from 2 weeks to 9 months. *AdipoR1* knockout mice were described in Rice et al [4]. Access to food and water was provided to animals *ad libitum*.

### In vivo Imaging

Before imaging, pupils were dilated with 1.0% cyclopentolate and 2.5 % phenylephrine (Akorn, IL). A 0.5% solution of proparacaine (Akorn, IL) was applied for corneal anesthesia. Mice were anesthetized by intraperitoneal injection of ketamine/xylazine (80.0 mg/kg /8.0 mg/kg). GenTeal® tears (Alcon, TX) were applied for corneal hydration. Fundus images were captured using a Micron III imaging system (Phoenix Research Labs, CA). A Bioptigen Envisu R2210 OCT system (Leica Microsystems, NC) was used to acquire a 1.8 mm long retinal scan (10 b-scans) centered over the optic nerve in an inferior-superior (vertical) and nasal-temporal (horizontal) orientation. B-scans were averaged, and retinal layers manually segmented using custom MATLAB code (Mathworks, MA). For each OCT data point, vertical and horizontal OCT scans were averaged for both eyes, and the central 200 μm over the optic nerve was excluded due to variable thickness. Infrared and autofluorescence images were obtained with a Spectralis HRA2 scanning laser ophthalmoscope (Heidelberg Engineering, MA) with a 55° lens.

### Histology

Immunohistochemical staining for ADIPOR1 and MFRP was conducted on tissues that were fixed in formalin and embedded in paraffin. The antibodies were rabbit anti-Adiponectin Receptor 1 (IBL, Catalog No. 18993, Japan) and goat anti-mouse MFRP (R&D Systems, Catalog No. BAF3445, Minnesota) respectively. Furthermore, H&E staining, RBP3 immunofluorescence staining, and AdipoR1 RNAScope assays were carried out on 5 μm thick flash-frozen sections. The primary antibody for RBP3 (also known as IRBP) was rabbit anti-RBP3 polyclonal (Proteintech, Catalog No. 14352-1-AP, Illinois). The *AdipoR1* RNAScope fluorescent assay was executed using an Automated Leica Bond RX, following the guidelines provided by the manufacturer. Specific probes for *AdipoR1* (Catalog No. 452858) and the RNAscope LS Multiplex Fluorescent reagent kit (Catalog No. 322800) were obtained from Advanced Cell Diagnostics (ACD, Newark, California, USA).

### RNA sequencing

Eye tissue was suspended in Buffer RLT/eye (1 eye/mouse; 10-18 mice/genotype) and prepared using the RNeasy kit (Qiagen, Germantown, MD). RNA quality was assessed by the 18S/28S rRNA bands in capillary electrophoresis (Bioanalyzer, Agilent Technologies, Santa Clara, CA). RNA sequencing was performed as described previously. Raw reads were aligned to the GENCODE mouse reference genome (GRCm38.p4) using STAR (v2.1.4a), with default settings. Read counts and transcripts per kilobase million (TPM) were calculated using RSEM (v1.2.22), and annotations of GRCm38.p4. On average, 10.8 million reads per sample were aligned to coding regions in mouse reference genome. Principal component analysis was performed to assess intra group and inter-group sample variations. Differential expression analysis was analyzed using the edgeR and limma packages. All data processing and analyses were performed in custom python and R scripts. Raw fastq files were deposited in Gene Expression Omnibus (GEO).

### Western blots

Method and general reagents for Western blots were described previously with a few modifications [10] Adipor1 and Tubulin antibodies utilized were rabbit anti-Adiponectin Receptor 1 (IBL, Catalog No. 18993, Japan) and rabbit anti-α Tubulin (Cell signaling, Catalog No. 2125, MA), respectively.

### Data Analysis

Quantification of protein levels were performed using GraphPad Prism Version 7 (GraphPad Software, San Diego, CA). OCT data were presented as mean± SD, with remaining data presented as mean ± SEM. Quantification or mRNA levels were performed using GraphPad Prism Version 10. Statistical differences were performed using one way ANOVA with Dunnett’s posttest. Differences were considered statistically significant for p values ≤ 0.05.

### Retinoid and DHA docking in ADIPOR1 crystal structure

*In silico* molecular docking was performed in both open and closed forms of ADIPOR1 monomers (Protein Data Bank (PDB) entry 6KRZ [6] using Schrödinger Maestro (Life Science: Maestro – Schrödinger, schrodinger.com). Default Protein Preparation Workflow parameters were used to add H atoms, fill in missing sidechains, and set residue protonation states at pH 7.4. As 6KRZ is a D208A mutant form of ADIPOR1, the open and closed monomers were converted back to wild type ADIPOR1 in Schrödinger Maestro prior to protein preparation. Docking receptor grids were generated around the center of mass of the co-crystallized ligands of the wild-type form of 6KRZ. 11-cis-retinaldehyde and DHA were docked using SP Glide [7] with default parameters.

## Results

Several labs have published the ocular phenotype in *Cfh^−/−^* mice [8]. Generally, the retinal phenotype is very mild and a subtle thinning in the photoreceptor layer was reported in mice almost 2 years of age [8, 9]. Our group obtained genetically modified *Cfh* mice from Dr. Marina Botto’s lab at Imperial College, UK. The mice were re-derived at Taconic Labs using C57BL/6Ntac mice, and then subsequently crossed to C57BL/6JbomTac animals to generate *Crb1 rd8* wild type mice. The eye phenotype observed in this line was in alignment with the modest eye pathology identified by others. The line was transferred to our *Cfh^+/−^* ; these were subsequently intercrossed to generate study cohorts and littermate controls. *Cfh^+/+^*, *Cfh^+/−^*, and 50% of *Cfh^−/−^* mice exhibited a normal fundus and OCT, as shown in figure 1A and B, respectively. However, the remainder of *Cfh^−/−^* mice examined (50%) exhibited a flecked retina and retinal degeneration, as shown in figure 1A and B, respectively. Quantification of OCT images revealed a reduction in total retinal thickness of approximately 33% in this subset of *Cfh^−/−^* mice (Figure 1C). Seven-month-old mice displayed a reduction in the number of photoreceptors in a subset of C*fh^−/−^* mice (Figure 1 D). The flecked retina and early onset retinal degeneration in a subset of *Cfh^−/−^*mice phenocopies that described in *AdipoR1^−/−^* deficient mice. *Cfh* and *AdipoR1* are in close proximity on mouse Chr 1. To test whether AdipoR1 contributes to the unexpected phenotype in *Cfh* mice, we set up an allele complementation test using matings between *Cfh^−/−^* and *AdipoR1^−/−^* mice with retinal degeneration (Figure 2 A and B, respectively) compared to an *AdipoR1^+/+^*control (Figure 2C). The mice from these matings are genetically double heterozygous for *AdipoR1* and *Cfh.* All of the offspring (*n*=13) exhibited retinal degeneration (Figure 2D), which demonstrates that a spontaneous mutation likely occurred in the AdipoR1 locus.

**Figure 1.**
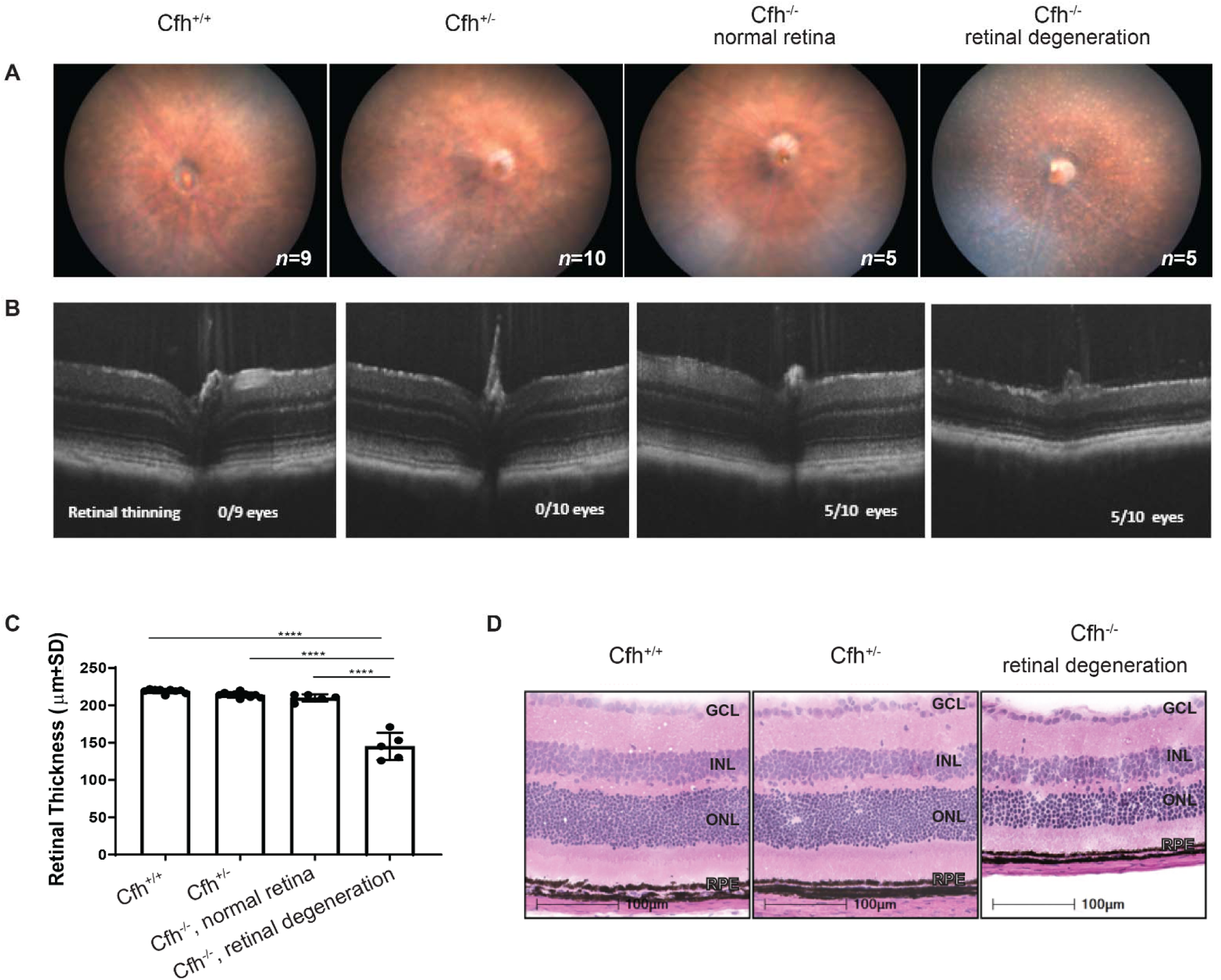
Retinal degeneration in a subset of *Cfh^−/−^* mice. (**A**) Fundus photography of 7-month-old *Cfh^+/+^* (*n* = 9), *Cfh^+/−^*(*n* = 10) and *Cfh^−/−^* (*n* = 10). The fundus in *Cfh^+/+^*and *Cfh^+/−^* mice appeared similar and were normal. Two phenotypes were observed in *Cfh^−/−^* mice. A subset (50%) exhibited no discernable phenotype and appeared similar to *Cfh^+/+^*and *Cfh^+/−^*. The remainder (50%) displayed hyper-reflective foci and retinal degeneration. (**B**) Optical coherence tomography (OCT) images of 7-month-old mice. (**C**) Total retinal thickness was measured from the basal RPE to the inner limiting membrane and represented averages of vertical and horizontal scans. A subset of *Cfh^−/−^* mice exhibited a 34% reduction in total retinal thickness attributable to PR loss. Images were of (**D**) Hematoxylin and eosin staining of cryosections from 4-week-old *Cfh^+/+^*, *Cfh^+/−^*, *Cfh^−/−^* mice. Photoreceptor loss was in a subset of 4-week-old *Cfh^−/−^* animals. GCL: ganglion cell layer; INL: inner nuclear layer; ONL: outer nuclear layer.

**Figure 2.**
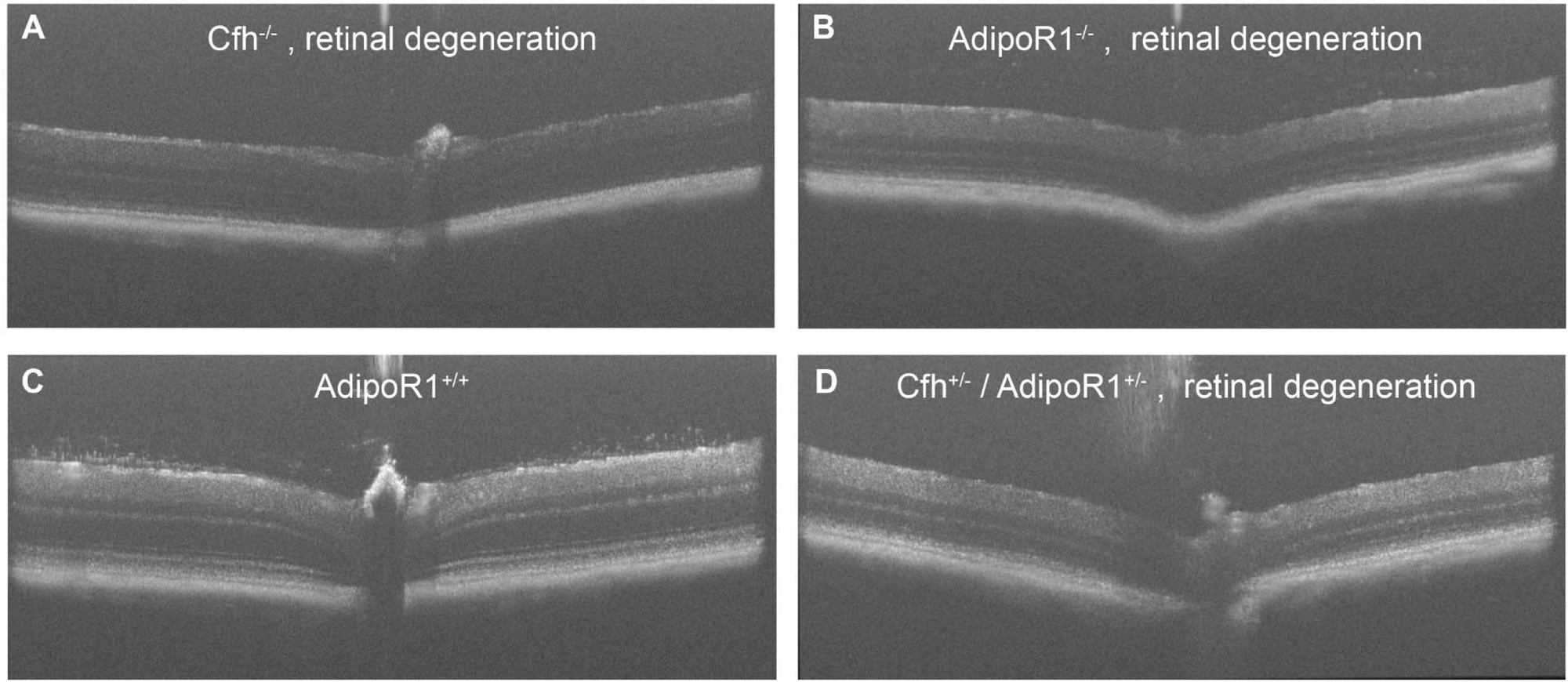
Complementation cross confirms AdipoR1 mutation is associated with retinal degeneration in *Cfh* mice. All double heterozygous offspring of a cross between a *Cfh^−/−^* mouse and an *AdipoR1^−/−^*mouse exhibit retinal degeneration, **A** and **B**, respectively. All offspring, *Cfh^+/−^ / AdipoR1^+/−^,* exhibited retinal thinning (D, *n =* 13). Image C is a representative OCT image of an *AdipoR1^+/+^* mouse.

Analysis of *AdipoR1* reads from our RNAseq data identified a missense mutation, c.841 C > T, in *AdipoR1* in *Cfh^−/−^* mice exhibiting retinal degeneration. The c.841 C > T mutation is 100% concordant in the *Cfh^−/−^* retinal degeneration samples (Figure 3). In contrast, about 50% of the *Cfh^+/−^* mice that did not show retinal degeneration were heterozygous for this point mutation. Intriguingly, one *Cfh^+/+^* and two *Cfh^+/−^* mice carried the c.841 C > T mutation in both alleles. These three mice, regardless of being WT or HET for the *Cfh* gene, exhibited retinal degeneration due to recombination between *Cfh* and *AdipoR1* (data not shown). This finding firmly establishes that retinal degeneration is caused by *AdipoR1* mutation rather than the *Cfh* genotype. The corresponding amino acid at position 281 is proline in the wild-type allele. A C to T mutation results in a conversion to serine (P281S) in ADIPOR1. This position is located within the V transmembrane domain of ADIPOR1.

**Figure 3.**
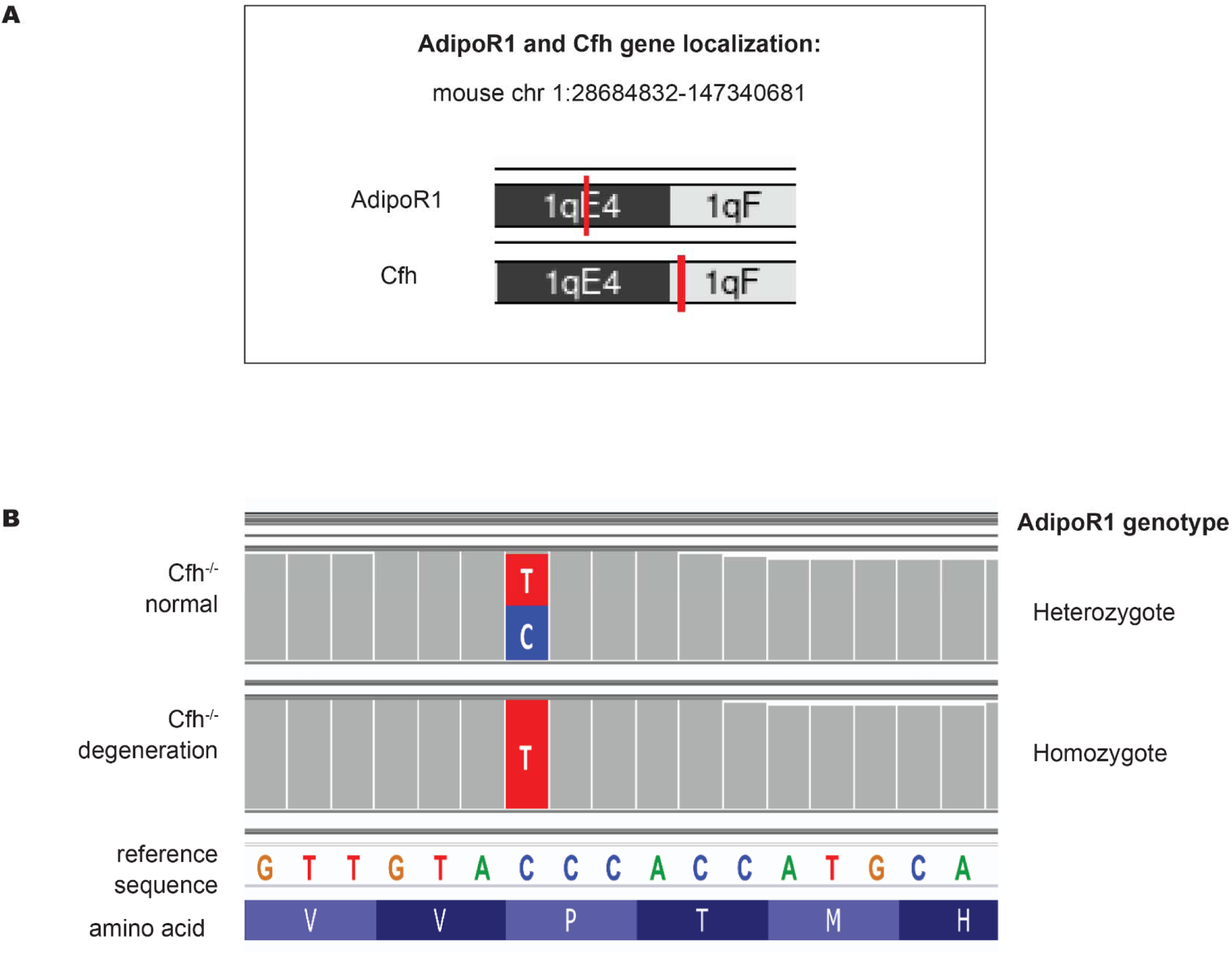
A missense mutation in the *AdipoR1* gene identified in Cfh mice with retinal degeneration. (**A**) *AdipoR1* and *Cfh* genes are located in close proximity on mouse Chr1. (**B**) Representative sequences of *AdipoR1* in two populations of *Cfh^−/−^*mice are shown. In *Cfh^−/−^* sequences with a normal retina, there is a cytosine to thymidine mutation on one allele. The other allele matches the wildtype reference sequence in the *AdipoR1* gene. In *Cfh^−/−^*mice with retinal degeneration, the point mutation is homozygous. Upon further breeding of the colony, the mutation was also found in one *Cfh^+/+^*and two *Cfh^+/−^* mice, which also exhibited retinal degeneration. Recombination between *AdipoR1* and *Cfh* genes explains this observation. The corresponding amino acid in the position is proline. The C to T mutation results in a proline to serine conversion at position 281. This confirms that the retinopathy is caused by the mutation in *AdipoR1,* regardless of the *Cfh* genotype.

Immunoanalysis using a validated, commercially available ADIPOR1 antibody [10] and *in situ* hybridization were employed to determine the impact of this point mutation on AdipoR1 protein and mRNA. The highest level of AdipoR1 immunoreactivity was observed in the apical microvilli of RPE cells (arrow in Figure 4 A). *Cfh^+/−^* mice with normal retinas exhibit ADIPOR1 expression (Figure 4 B), albeit at various levels likely due to heterozygosity of the c.841 C > T mutation. ADIPOR1 staining is manifestly lower in *Cfh^−/−^* mice with retinal degeneration (Figure 4 C). We asked if the absence of ADIPOR1 could be due to altered or absent apical microvilli, but staining with MFRP showed enrichment in apical microvilli, indicating that the structures were still present regardless of Cfh genotype (Figure 4 D, 4 E and 4 F). This result is comparable to a prior report describing *AdipoR1* gene targeted knockout mice [10]. Western blot confirmed decreased levels of ADIPOR1 protein in *Cfh^−/−^* eyes with retinal degeneration (Figure 5 A). Densitometry revealed a 75% reduction of ADIPOR1 in *Cfh^−/−^* eyes with retinal degeneration (Figure 5 B). Finally, *in situ* hybridization revealed that *AdipoR1* signal is comparable in neural retina and RPE cells in *Cfh^+/+^ Cfh^+/−^* and *Cfh^−/−^* mice with retinal degeneration (Figure 6). This result between *AdipoR1* transcript and protein levels is not technical as the RNAscope probe set contains 20 pairs of probes binding to different regions of the mRNA sequence. The presence of wild-type levels of AdipoR1 transcript but reduced ADIPOR1 protein in *Cfi^−/−^* mice strongly suggests that the point mutation affects ADIPOR1 protein production, stability, or half-life. ADIPOR1 is required for normal retinal visual processing and health [5, 10–12].

**Figure 4.**
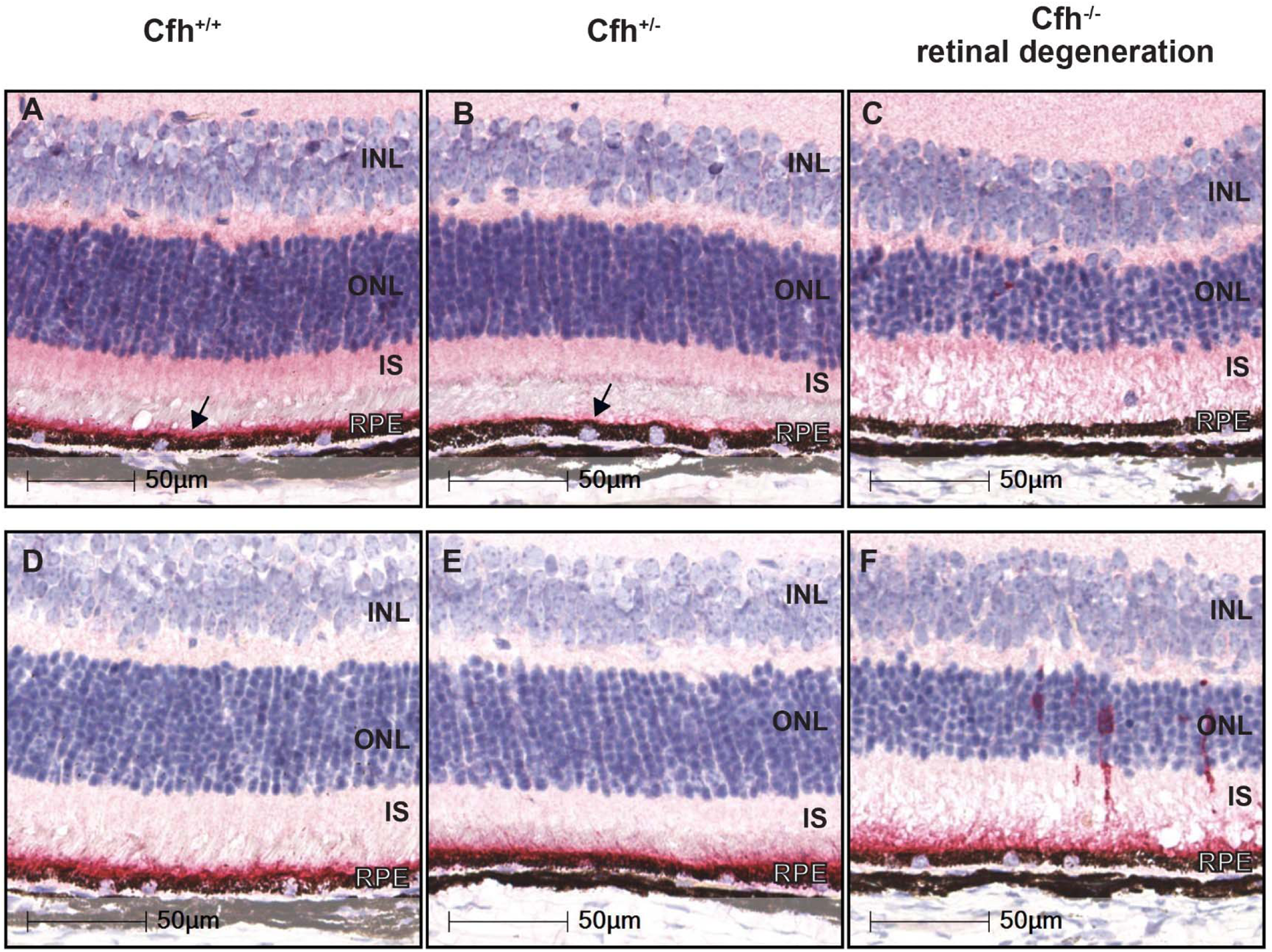
Immunostaining of ADIPOR1 and MFRP in *Cfh* mice. ADIPOR1 (red stain, arrow) is highly expressed in RPE and their apical microvilli in *Cfh^+/+^* (**A**) and in *Cfh^+/−^* (**B**) mice that exhibit a normal retina, albeit with reduced intensity in this *Cfh^+/−^* example (likely a heterozygous for the c.841 C > T mutation). ADIPOR1 levels are dramatically decreased in *Cfh^−/−^* with retinal degeneration (**C**). Membrane Frizzled-Related Protein (MFRP) is expressed in the RPE apical microvilli, regardless of genotype *Cfh^+/+^* (**D**), *Cfh^+/−^*(**E**) and retinal degeneration in *Cfh^−/−^* (**F**), demonstrating that ADIPOR1 deficiency is not due to absence of RPE microvilli.

**Figure 5.**
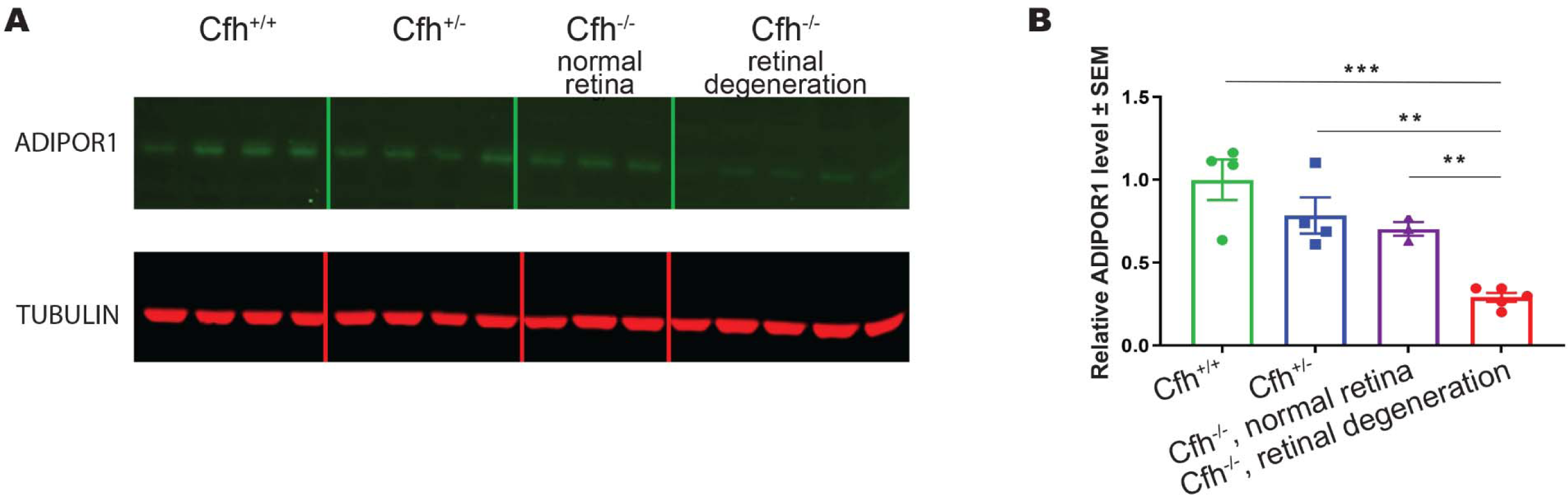
Western blots of ADIPOR1 in *Cfh^+/+^, Cfh^+/−^, Cfh^−/−^* with normal retinal anatomy compared to *Cfh^−/−^* with retinal degeneration. *Cfh^−/−^* mice with retinal degeneration exhibit reduced levels of ADIPOR1 expression compared to *Cfh^+/+^, Cfh^+/−^ and Cfh^−/−^* cohort littermates with normal retinas. Tubulin was used as a loading control. **B.** Densitometry quantification of western blots demonstrate about 75% reduction in ADIPOR1 protein in the retinal degeneration samples (** p< 0.01; ***p<0.001).

**Figure 6.**
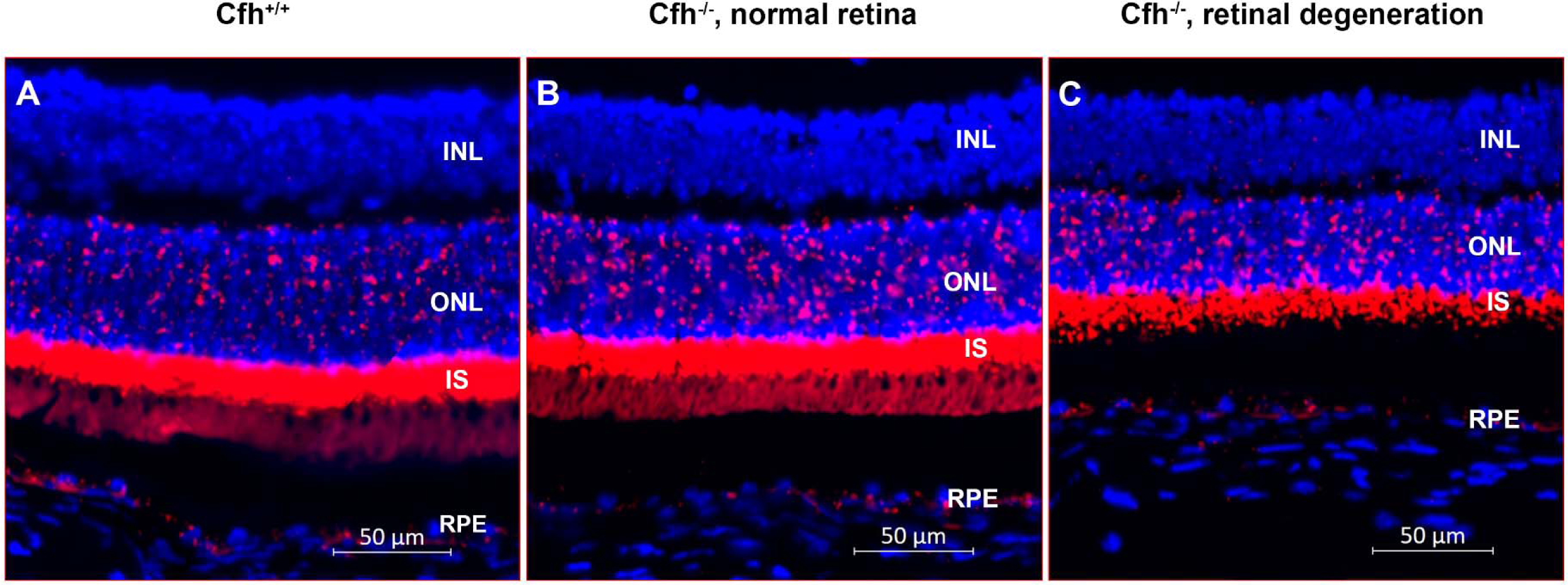
*AdipoR1* mRNA is detected in all Cfh genotypes examined, regardless of retinal anatomy. An *AdipoR1*-specific riboprobe (red) is located throughout the neural retina and RPE in both *Cfh^+/+^*(**A**) and *Cfh^−/−^* mice with no degeneration (**B**). *AdiopR1* is also present in *Cfh^−/−^* mice with retinal degeneration (**C**).

We further characterized molecular and anatomical changes in postnatal day (PD)15, and PD30 *AdipoR1* knockout mice to understand early functional consequences of ADIPOR1 deficiency. Bulk RNAseq revealed a significant upregulation of retinol binding protein 3 (*Rbp*3) in PD15 *AdipoR1^−/−^*mice, while the *AdipoR1^+/−^* and *AdipoR1^+/+^* mice were comparable (Figure 7 A). *Rbp3*, also known as interphotoreceptor retinol binding protein (IRBP), is expressed by photoreceptors, and secreted into the interphotoreceptor matrix (IPM) where it shuttles retinoids between the RPE and photoreceptors. Another IPM gene, Interphotoreceptor matrix proteoglycan 2 (*Impg2*), was not different among the genotypes (Figure 7B), suggesting that the IPM is not globally disrupted. IRBP staining in PD30 *AdipoR1^+/+^* mice appears as two distinct bands in the IPM (Figure 7 C). In contrast, IRBP is broadly distributed throughout the IPM in *AdipoR1^−/−^* mice (Figure 7 D).

**Figure 7.**
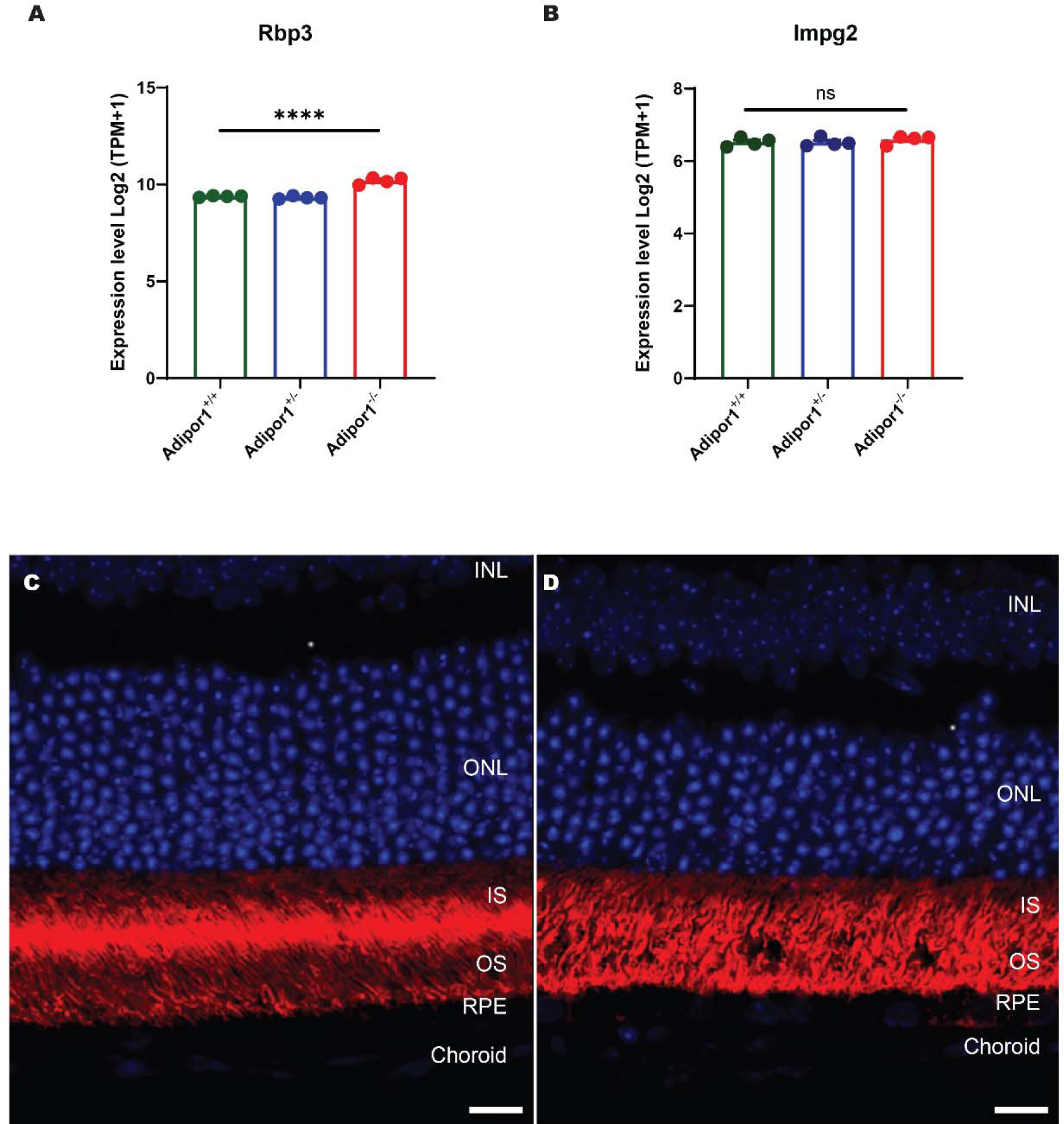
Abnormal levels of *Rbp3* mRNA and altered spatial distribution of RBP3 protein. (**A**) *Rbp3* mRNA is significantly upregulated in postnatal day (PD) 15 *AdipoR1^−/−^* mice (KO) by RNAseq compared to *AdipoR1^+/−^* (HET) and *AdipoR1^+/+^* (WT). *p* < 0.0001. Another interphotoreceptor matrix gene, *Impg2*, is similar among the genotypes. *p* = 0.4614. (**C**) RBP3 (aka IRBP) protein in PD30 *AdipoR1^+/+^* is observed in two bands, one in the inner segments (IS) and the other at the tip of the OS, bordered by the outer nuclear layer (ONL) and (RPE). (**D**) In contrast, RBP3 is broadly distributed in the is and os region in *Adipor1^−/−^* mice (*n* = 3 per genotype). Scale bar is 10 μm for **C** and **D.**

A recently published crystal structure of ADIPOR1 (PDB entry 6KRZ) was selected for molecular modeling [6]. 6KRZ consists of an asymmetric unit of three ADIPOR1 monomers, two of which contain ligands co-crystalized in the transmembrane pore (Figure 8A). Examination of the monomers reveals that they can assume either closed or open forms (Figure 8B). In the closed form, the intracellular end of helix V is closer to the other transmembrane helices, causing the transmembrane pore to be narrower and tube-like. In the open form, the intracellular end of helix V moves away from the other transmembrane helices, causing the transmembrane pore to be wider and channel-like. Both closed and open forms are seen with co-crystalized ligands in 6KRZ. No structural changes were observed in the intracellular domain between the open and closed forms. 6KRZ contains a single-point D208A mutation which was changed to wild type ADIPOR1 prior to modeling. Conversion of Alanine 208 back to the wild-type Aspartate residue did not produce noticeable changes in the ADIPOR1 tertiary structure.

**Figure 8.**
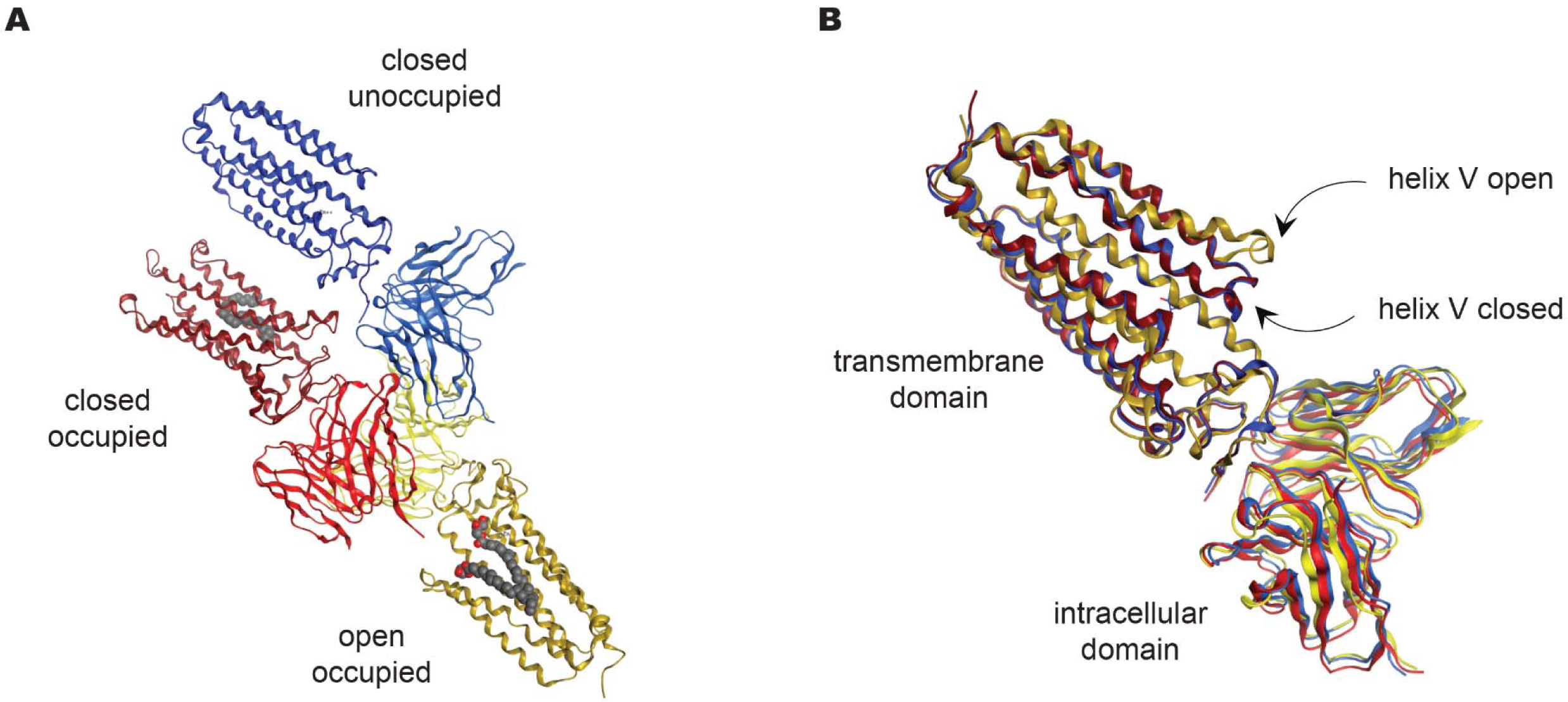
Crystal structure of ADIPOR1 (PDB entry 6KRZ) comprised of three monomers. (**A**) The individual monomers display different combinations of helix V position and ligand occupancy: closed/occupied (red), closed/unoccupied (blue), open/occupied (yellow). (**B**) Alignment of the monomers illustrates the similarities in the intracellular domain and the differences in helix V in the transmembrane domain. Co-crystalized ligands have been removed for clarity.

*In silico* docking of 11-cis-retinaldehyde and DHA was performed using the closed and open forms of wild type ADIPOR1 (Figure 9). Despite the differences in pocket shape resulting from the position of helix V, the ligands docked into both the closed and open forms of ADIPOR1 in similar manners (11-cis-retinaldehyde in Figure 9A and 9B; DHA in Figure 9C and 9D). The aldehyde oxygen of 11-cis-retinaldehyde and the carboxylic acid of DHA both coordinate with the Zn^2+^ ion present in the transmembrane pore, while the carbohydrate tails of the ligands bind nonspecifically to the rest of the lipophilic pocket. The docking results show that both open and closed states of ADIPOR1 can accommodate 11-cis-retinaldehyde and DHA, suggesting that ADIPOR1 may be involved in trafficking retinoids and fatty acids.

**Figure 9.**
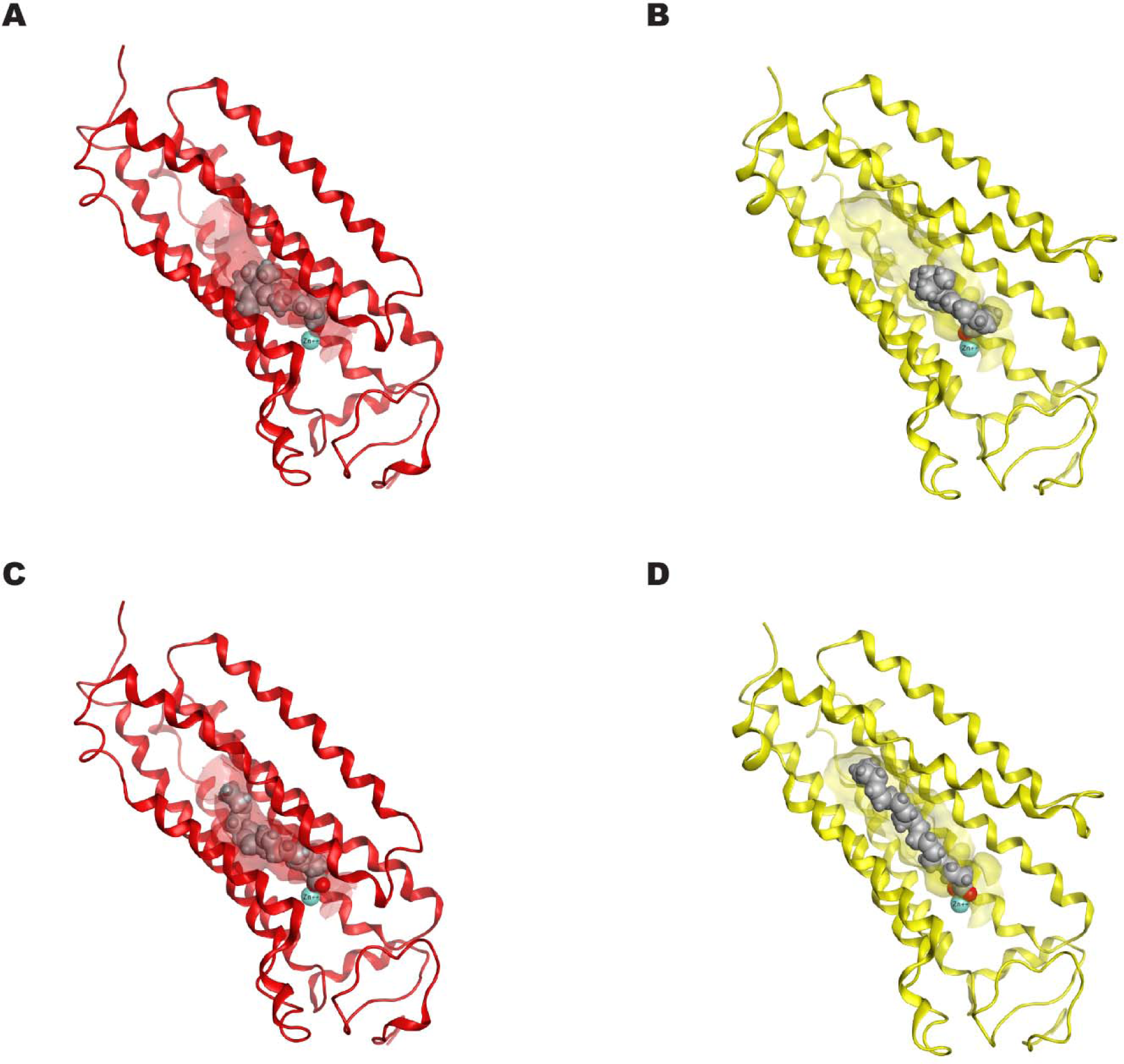
Molecular docking poses of 11-cis-retinaldehyde in (**A**) closed form and (**B**) open form structures of ADIPOR1. Molecular docking poses of DHA in (**C**) closed form and (**D**) open form structures of ADIPOR1. The oxygen atoms of both ligands coordinate with the Zn^2+^ ions (blue spheres) in the pocket. In the helix V closed state docking (red structures) the binding pocket is tube-like in shape (faint red), while in the helix V open state docking (yellow structures) the binding pocket is solvent exposed on the helix V side and assumes a channel-like shape (faint yellow).

## Discussion

An important outcome of this study is to further highlight awareness of spontaneous mutations that arise in research colonies. These serendipitous findings open the door for new research projects [13]. The retinal pathology discovered in our *Cfh* knockout colony is identical to the phenotype of *AdipoR1* knockout mice [4]. Given the similarity of this ocular phenotype to that of *AdipoR1*-deficient mice, and the fact that *Cfh* and *AdipoR1* genes are located near each other on chromosome 1, suggested a mutation in *AdipoR1* is responsible for the phenotype. This work led us to the discovery of a missense mutation in the *AdipoR1* gene. This allele decreases ADIPOR1 but residual protein is observed at about 25% of wild type levels. Discovery of this allele may facilitate future studies of ADIPOR1 structure / function and provide cellular assays for ligand interactions and receptor activity.

The crystal structures of ADIPOR1 and ADIPOR2 (a structurally similar receptor) have been solved and their transmembrane pores contain a mono-unsaturated fatty acid, oleic acid. ADIPOR1 features a seven transmembrane (7TM) structure with substantial transmembrane pores and a zinc-binding site near the intracellular surface within the 7TM domains [6, 14]. The zinc ion at this site is coordinated by three histidine residues, with the histidine and aspartate residues in ADIPOR1 being highly conserved across a range of species. Two labs have demonstrated that ADIPOR1 has ceramidase activity [11, 14]. *AdipoR1^−/−^*mice accumulate ceramide species in both the retina and the RPE. Increased levels of ceramide lead to photoreceptor cell death, which can be attenuated with low molecular weight inhibitors of ceramide production [11]. ADIPOR1 may have hydrolase activity, with the zinc-binding site potentially acting in the catalytic domain. The large cavity within ADIPOR1 opens significantly towards the intracellular surface and another small opening towards the outer lipid layer’s middle, possibly functioning as pathways for substrates and products of the hydrolytic activity. Human ADIPOR1 can assume two closed and one open structures, likely in equilibrium, with transitions between these states possibly linked to a signaling mechanism [6]. The ADIPOR1 open conformation renders the putative catalytic site and substrate binding domain exposed to the cytoplasm and fully accessible to the inner membrane leaflet.

The c.841 C > T mutation in the *AdipoR1* gene led to the substitution of proline with serine at position 281 in the ADIPOR1 protein sequence. The proline at position 281 (P281) is evolutionarily conserved and located in the fifth transmembrane (TM) domain, near the extracellular side of the cell membrane. The P281 residue plays a critical role in the conformational dynamics of ADIPOR1, especially in the open and closed positions of the fifth TM domain [6]. In the closed conformation, P281 contributes to the inward bending of the fifth TM, while in the open conformation, it aids in maintaining a straighter structure. Therefore, the mutation from the nonpolar proline to the polar serine at this position is hypothesized to adversely affect the normal conformation and function of ADIPOR1.

ADIPOR1 has been shown to be required for accumulation of docosahexaenoic acid (DHA) in the retina, which can be elongated into very long-chain fatty acids that are important for retinal health [4, 15]. Long-chain polyunsaturated fatty acids are enriched in photoreceptor outer segment disk phospholipids providing membrane fluidity that supports the visual cycle cascade. The critical role of ADIPOR1 in vision biology is underscored by observations that *AdipoR1* knockout mice models [4,10] develop retinal degeneration and recent discoveries of two novel human mutations in ADIPOR1 leading to retinitis pigmentosa [4, 5, 12]. One of the earliest physiological effects of ADIPOR1 deficiency in mice is a dramatic reduction of photopic a and b wave electroretinograms (ERGs), which are severely diminished as early as 3 and 4 weeks of age [11, 15]. Loss of photoreceptors is apparent in histological samples of *AdipoR1^−/−^* mice at 3 and 4 weeks of age, but the magnitude of loss does not support the profound effect observed in the ERGs [11, 15]. RHODOPSIN is detectable by immunohistochemistry in rod outer segments at 3 weeks of age in *AdipoR1^−/−^* mice. However, RHODOPSIN levels are significantly lower as revealed with Western blot analysis [10, 15]. Ultrastructure analysis at 2 weeks of age revealed disorganized outer segments in *AdipoR1^−/−^* mice [15].

*AdipoR1^−/−^* mice have a dramatic reduction of 11-cis-retinaldehyde as early as 1 month of age when compared to wild type animals [11]. In addition, there is a significant elevation of IRBP mRNA and protein as early as PD15 [10]. The spatial localization of IRBP is also abnormal in the *AdipoR1* KO mice. IRBP facilitates trafficking of 11-cis-retinal released from the RPE in the interphotoreceptor matrix (IPM). The upregulation of IRBP could be a compensatory mechanism in the presence of dramatic reductions in the levels of 11-cis-retinal.

As mentioned above, mice deficient in ADIPOR1 display several phenotypes such as massive reductions in 11-cis-retinaldehyde at a young age and reduced ERG responses in dark-adapted conditions, indicative of disruptions in the retinoid cycle. In addition, ADIPOR1 mice exhibit lowered levels of DHA-containing phospholipids in the retina, and disorganized outer segments as early as two weeks of age. Knockout of other genes reproduces some of the features of ADIPOR1 deficient mice. Deficiency in the DHA transporter Mfsd2a leads to a reduction in DHA levels that is similar in magnitude to that observed in ADIPOR1 deficient mice [16, 17]. However, *Mfsd2a* KO mice exhibit slow photoreceptor loss and shortening of the OS, but no significant abnormalities in the overall ultrastructure of the OS or the spacing of the OS discs and their visual signal transduction remains unaffected. Additionally, there is no reduction in the expression and localization of phototransduction protein components as observed in *AdipoR1^−/−^* mice [10]. These results suggest that decreased DHA levels alone do not explain the unique phenotype in the ADIPOR1 mutant animals. Deficiency of the retinoid isomerohydrolase RPE65 leads to subtle and gradual retinal photoreceptor degeneration, complete loss of rod function measured with ERGs, and a complete loss of rhodopsin despite presence of outer segments, albeit with disk abnormalities as viewed with electron microscopy [18]. The electroretinogram responses of *Rpe65^−/−^* mice in dark-adapted conditions are severely diminished due to the lack of 11-cis-retinaldehyde. Mice deficient in Lecithin-retinol acyltransferase (LRAT) exhibit dramatic decreases in RHODOPSIN and 11-cis-retinaldehyde. The outer segment length is shortened and a slowly progressing rod photoreceptor degeneration that is comparable to the RPE65-deficient mice [19]. Therefore, the decrease in 11-cis retinaldehyde observed in ADIPOR1-deficient mice by itself does not explain the retinal degeneration phenotype.

Comparison of retina phenotypes observed in several animal models with retinoid cycle defects or DHA transport disorders suggests that ADIPOR1 is involved in both DHA transport and the retinoid cycle. IRBP binds both DHA and retinoids and seems uniquely upregulated in ADIPOR1 mice [20–23]. Enrichment of transmembrane ADIPOR1 in apical microvilli of RPE cells places ADIPOR1 in the region where retinoids and DHA cross the membrane to traffic to IRBP in the IPM. ADIPOR1 lacks G-protein activity and has an inverted topology in the membrane. The N-terminus is intracellular, and the C-terminus is facing the IPM. Additional studies will determine if ADIPOR1 interacts with retinoid carrier proteins such cellular retinaldehyde binding protein (CRALBP), cellular retinoid binding protein (CRBP) and/or IRBP (reviewed in [24] to facilitate the efficient transport of retinoids and DHA between RPE and photoreceptors. The data suggests that deficiency in both 11-cis-retinadehyde and DHA levels, combined with elevated levels of ceramide species in the RPE contributes to the unique phenotype in mice lacking ADIPOR1.

## Acknowledgements

The authors would like to thank Chris Wilson for their thoughtful review of the manuscript, Sha-Mei Liao, Tom Vollmer and Siyuan Shen for their support on the project.

